# Transfer of antigen-encoding bone marrow under immune-preserving conditions deletes mature antigen-specific B cells in recipients and inhibits antigen-specific antibody production

**DOI:** 10.1101/2019.12.20.885343

**Authors:** Jeremy F. Brooks, Janet M. Davies, James W. Wells, Raymond J. Steptoe

**Author notes:** Corresponding author: R. J. Steptoe, UQ Diamantina Institute, Level 6, Translational Research Institute, 37 Kent Street, Woolloongabba, Queensland, Australia 4102.

## Abstract

Pathological activation and collaboration of T and B cells underlies pathogenic autoantibody responses. Existing treatments for autoimmune disease cause non-specific immunosuppression and induction of antigen-specific tolerance remains an elusive goal. Many immunotherapies aim to manipulate the T-cell component of T-B interplay but few directly target B cells. One possible means to specifically target B cells is the transfer of gene-engineered BM that, once engrafted, gives rise to widespread specific and tolerogenic antigen expression within the hematopoietic system. Gene-engineered bone marrow encoding ubiquitous ovalbumin expression was transferred after low-dose (300cGy) immune-preserving irradiation. B-cell responsiveness was monitored by analyzing ovalbumin-specific antibody production after immunization with ovalbumin/complete Freund’s adjuvant. Ovalbumin-specific B cells and their response to immunization were analyzed using multi-tetramer staining. When antigen-encoding bone marrow was transferred under immune-preserving conditions, cognate antigen-specific B cells were purged from the recipient’s pre-existing B cell repertoire as well as the repertoire that arose after bone marrow transfer. OVA-specific B-cell deletion was apparent within the established host B-cell repertoire as well as that developing after gene-engineered bone marrow transfer. OVA-specific antibody production was substantially inhibited by transfer of OVA-encoding BM and activation of OVA-specific B cells, germinal centre formation and subsequent OVA-specific plasmablast differentiation were all inhibited. Low levels of gene-engineered bone marrow chimerism were sufficient to limit antigen-specific antibody production. These data show that antigen-specific B cells within an established B-cell repertoire are susceptible to *de novo* tolerance induction and this can be achieved by transfer of gene-engineered bone marrow. This adds further dimensions to the utility of antigen-encoding bone marrow transfer as an immunotherapeutic tool.

## Introduction

Pathogenic antibodies arising through failure of immune tolerance mechanisms contribute to a range of clinically-important conditions. Pathogenic autoantibodies mediate immune complex deposition and complement-dependent tissue damage (e.g. Hashimoto’s disease, Goodpasture’s syndrome/rheumatoid arthritis), activate or block hormone or other receptors (e.g. Grave’s disease, Myasthenia gravis), promote self-antigen presentation (e.g. type 1 diabetes) or inhibit the activity of protein replacement therapies (e.g. Factor VIII) (Suurmond and Diamond 2015). In allergy, allergen-specific IgE precipitates acute allergic reactions and promotes the ‘atopic march’ and the perpetuation of allergic disease. To alleviate the effects of antibody-mediated pathology most therapeutic approaches employ non-specific immune-suppression to target the consequences of antibody-mediated effects, rather than controlling the immunological root cause of disease. Ultimately, therapies that prevent development and ongoing production of pathogenic antibodies by reinstating immune tolerance or that strip the repertoire of problematic B cell and antibody specificities are required. Because of their capacity to antigen-specifically inhibit the immunological mechanisms driving pathogenic B-cell activation and antibody production, antigen-specific immunotherapies are looked to as promising future approaches.

Because they are an ongoing source of antibody production and resistant to many interventions (Hiepe and Radbruch 2016) long-lived plasma cells have traditionally been perceived as a key challenge for attempts to modulate the homeostasis of established pathogenic antibody responses. Less appreciated, perhaps, is the importance of short-lived plasma cells and, particularly, antigen-specific plasmablasts. These are short-lived and arise rapidly in large numbers after B-cell activation, but are an important, major source of pathogenic antibodies in some autoimmune diseases and potentially in allergy (Teng, Wheater et al. 2012, Kerkman, Rombouts et al. 2013, Wu and Scheerens 2014, Stathopoulos, Kumar et al. 2017, Hale, Rawlings et al. 2018). New drug (Neubert, Meister et al. 2008) and biologics–based (Ramanujam, Wang et al. 2006, Cogollo, Silva et al. 2015) approaches, may overcome the hurdle presented by long-lived plasma cells, so purging B cells with potentially problematic specificities or limiting their activation and/or differentiation to antibody-secreting cells (i.e. plasmablasts and plasma cells) represents what is likely a key goal of immunotherapies for antibody-mediated diseases in the future. Current immunotherapeutic approaches such as antigen-specific immunotherapy (SIT) and peptide immunotherapy aim to indirectly limit B cell responsiveness by modulating T cells, either to induce regulatory T cells or to divert antibody responses to ‘non-pathogenic’ isotypes (Sabatos-Peyton, Verhagen et al. 2010). A more effective approach might be to directly target B cells, either for deletion or inactivation, but how to achieve that using conventional immunotherapies remains undefined. One particular challenge for directly, and antigen-specifically, targeting B cells appears to be that the antigen delivery methods of conventional immunotherapies, while capable of modulating T cell responses, are unsuited to tolerogenic antigen delivery to B cells. However, success could alleviate pathogenic antibody-mediated inflammation in some seropositive diseases such as rheumatoid arthritis or the expansion of the effector T-cell response in T cell-mediated diseases such as type 1 diabetes or multiple sclerosis (Marino, Walters et al. 2014, Jelcic, Al Nimer et al. 2018, Getahun and Cambier 2019).

One emerging approach showing great promise for antigen-specific immunotherapy is hematopoietic stem cell (HSC)-mediated gene therapy. In this setting, transfer of bone marrow (BM) or HSC that carry a genetic construct encoding antigen(s) gives rise to progeny that express antigen and confer robust immune tolerance. When used with mild pre-transfer conditioning unwanted pathogenic responses can be antigen-specifically targeted while bystander immunity is preserved (Coleman, Bridge et al. 2013). For T cells, this approach has been shown to inactivate both naïve and memory responses and can override tolerance defects (Steptoe, Ritchie et al. 2003) and the effects of inflammation (AL-Kouba, Wilkinson et al. 2017). Early studies demonstrated that chronic ligation of the BCR by membrane-bound antigen was a powerful stimulus for induction of both central and peripheral B-cell tolerance (Goodnow, Crosbie et al. 1988, Hartley, Crosbie et al. 1991). Advantageously, for B-cell tolerance, transfer of gene-engineered BM or HSC can introduce permanent expression of an antigen of interest into sites of B-cell development, selection and peripheral residence. This, then provides a means of antigen delivery that overcomes the limitations of conventional immunotherapy for direct B-cell tolerogenesis.

Exploration of HSC-mediated gene therapy to target B-cell responses has been limited and, when studied, has typically used conditions where lethal irradiation and/or leukocyte depletion have been used (Baranyi, Linhart et al. 2008, Chung, Figgett et al. 2014). While this might recapitulate B-cell development as the immune system is reinstated after BM or HSC transfer, importantly such immunoablative approaches do not address one of the most critical questions for clinically-applicable immunotherapy. Which is “whether mature, established antigen-specific B cells can be specifically targeted and purged, for example by antigen-specific deletiuon, from within a normal homeostatic repertoire of B cells while leaving the remaining B-cell repertoire intact?” Here we address this and ask can ‘*de novo* peripheral tolerance’ be induced in a pre-established B-cell compartment? We use recent advances in BM/HSC-mediated gene therapy that use mild ‘conditionng’ regimes to enable the effects on established lymphocyte populations and their repertoires to be examined (Coleman, Bridge et al. 2013, Coleman, Jessup et al. 2016, Bhatt, Rudraraju et al. 2017). Many studies of B-cell tolerance employ systems where the B-cell receptor (BCR) is genetically manipulated either to increase clonal frequency.. But, tolerance processes in settings where BCR is genetically modified may not fully replicate those in a non-BCR manipulated setting. Therefore we employed newly-developed tracking tools that, for the first time in BM/HSC transfer studies of this kind, enable the detection and tracking of naïve antigen-specific B cells at the cellular level within a ‘normal’ polyclonal repertoire.

We now show that mature B cells specific for the antigen expressed *de-novo* after BM transfer are deleted from within the normal polyclonal repertoire,, in recipients. Consequently, antigen-specific B cell responses in recipients are limited and production of antigen-specific antibodies is inhibited in recipients. Transfer of gene-engineered BM/HSC therefore delivers antigen in a form that is directly tolerogenic to mature antigen-specific B cells in recipients and inhibits antibody production.

## Results

### Transfer of antigen-encoding BM under myeloablative conditions reveals an antigen abundance threshold for inhibition of antibody production

To test whether transfer of antigen-encoding BM induces B-cell tolerance and also to understand whether the prevalence of antigen-expressing cells dictates the outcome, we generated BM chimeras with graded proportions of OVA-expressing cells (Fig. 1A). For this, BM from act.OVA (CD45.2^+^) mice was mixed in increasing proportions with non-transgenic (non-Tg) BM (CD45.1^+^) and transferred to lethally-irradiated (1100cGy) recipients (Fig. 1A). Ten weeks post-transfer, BM had engrafted and repopulated the host such that the proportion of act.OVA leukocytes closely reflected the proportion within transferred BM (Fig. 1B). Overall, total spleen cellularity, the overall number of T cells, B cells and myeloid lineage cells in recipient mice reflected that in no-BMT controls and the proportion of act.OVA-derived T cells, B cells, myeloid cells and DC reflected that for overall chimerism.

**Figure 1.**
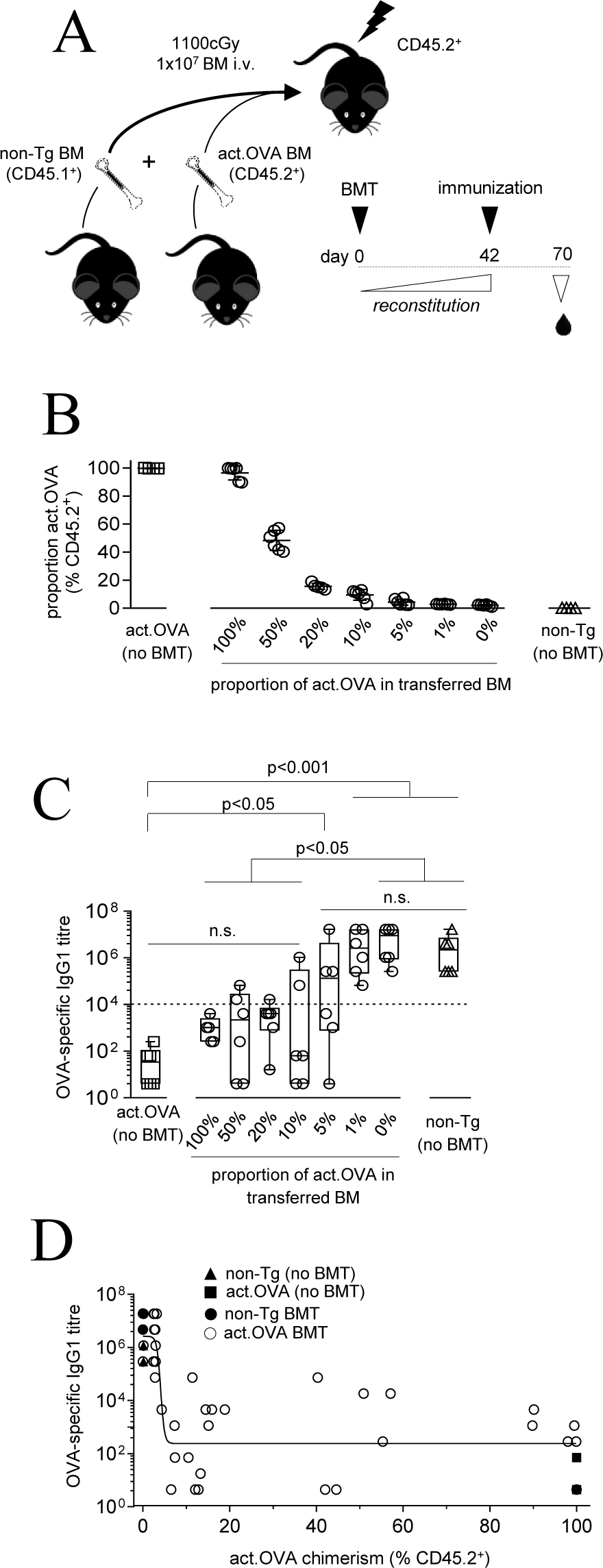
Antigen prevalence determines the extent of antigen-specific B cell tolerance. **A-D)** 10^7^ BM containing graded proportions of act.OVA BM (CD45.2^+^) and B6.SJL BM (CD45.1^+^) was transferred to lethally-irradiated (1100cGy) C57BL/6J (CD45.2^+^) recipients. Untransplanted controls (act.OVA and non-Tg C57BL/6J) were also used. **A)** experimental layout. 6 weeks after BM transfer, mice were immunized with OVA/CFA and analyzed 4 weeks later. **B)** act.OVA chimerism was determined as the proportion of CD45.2^+^ (act.OVA) cells in spleen 10 weeks after BM transfer, **C)** OVA-specific IgG1 titres 4 weeks following OVA/CFA immunization, **D)** relationship of OVA-specific IgG1 production to act.OVA chimerism. Plots show individual mice with mean±SD (B) or median and interquartile range (C). The dotted line indicates the antibody titre below which mice were considered ‘tolerant’. Data are pooled from 2 independent experiments. ANOVA with Tukey’s post-hoc test of log_10_-transformed titres (C) and log-inhibitor vs response curve (D).

To understand the impact of BM transfer on B-cell responsiveness, we immunized recipients and controls with OVA/CFA and 28 days later measured OVA-specific IgG1. Immunization induced a high level of OVA-specific IgG1 in non-Tg mice (titre typically >10^5^, Fig. 1C, Suppl. Fig. 1) whereas substantially lower levels of OVA-specific IgG1 (titre typically <10^3^, Fig. 1C, Suppl. Fig. 1) were induced in act.OVA mice. This was as expected as the OVA-specific antibody response in act.OVA mice after OVA/CFA immunization is transient and generates only low-affinity antibodies. Development of OVA-specific IgG1 showed an inverse relationship to the proportion of act.OVA BM transferred (Fig. 1C). When act.OVA cells comprised 10% or more of transferred BM, anti-OVA IgG1 titres, while variable, did not differ overall from the very low levels in act.OVA controls and were reduced relative to non-Tg no BMT controls or mice that received non-Tg BM only (0% act.OVA) (Fig. 1C, Suppl Fig. 1).

When transferred BM contained 5% act.OVA cells, OVA-specific IgG1 titres in recipients were variable but, mostly, intermediate between act.OVA and non-Tg no BMT controls (Fig. 1C). In contrast, after transfer of non-Tg BM or BM containing 1% act.OVA cells OVA-specific IgG1 titres were high and equivalent to those in non-transgenic controls (Fig. 1C). Plotting titre vs abundance of OVA^+^ leukocytes revealed that maximal inhibition of OVA-specific IgG1 was achieved when the proportion of act.OVA-derived leukocytes exceeded approximately 10% (Fig. 1D). These data indicate inhibition of the B-cell responsiveness that leads to antibody production occurs when relatively low proportion of leukocytes (~10%) express OVA.

### Transfer of antigen-encoding BM under mild, immune-preserving conditions limits antigen-specific antibody production

Low-dose irradiation serves as a mild conditioning regime that permits significant engraftment of transferred BM whilst substantially preserving recipient immunity (AL-Kouba, Wilkinson et al. 2017, Bhatt, Rudraraju et al. 2017). Transfer of BM encoding for antigen expression following low-dose irradiation is an effective means of ablating antigen-specific T-cell responsiveness (Coleman, Bridge et al. 2013). We tested whether such transfer of antigen-encoding BM would limit antigen-specific antibody responses. Graded doses of OVA-encoding BM was transferred to non-Tg recipients after low-dose (300cGy) irradiation (Fig. 2A). A short course of immunosuppression was used to prevent recipient immune-mediated rejection of OVA-expressing BM. Ten weeks after BM transfer, the emergence of donor-derived leukocytes was related to the dose of BM transferred (Fig. 2B). Total spleen cellularity, and the number of T cells, B cells, myeloid cells and DC did not differ across treatment groups. To probe for antibody production, we immunized recipients and controls with OVA/CFA and measured OVA-specific IgG1 antibodies in the blood. Recipients of non-Tg BM produced similar levels of OVA-specific IgG1 as non-Tg no BMT controls regardless of whether they were administered rapamycin or not (Fig. 2C, Suppl. Fig. 2) indicating the procedures themselves did not limit antibody production. Overall, OVA-specific IgG1 production was reduced by transfer of act.OVA BM (Fig 2C, D) and inhibition was greater as the dose of act.OVA BM used increased. Examining the extent of OVA-expressing leukocyte chimerism it was apparent that even low levels of chimerism were sufficient to modulate OVA-specific IgG1 production (Fig. 2C). This was reflected in the inhibition vs dose (OVA^+ve^ leukocyte abundance) curve which showed that levels of chimerism greater than approximately 15% reduced OVA-specific IgG1 titres by 2 to 4 orders of magnitude (Fig. 2D). As OVA-specific antibody production was reduced by such a large scale, but in most mice a large proportion of leukocytes remain recipient-derived after BMT (~40-80%, Fig. 2B), we conclude that the capacity to produce OVA-specific antibodies was purged from the mature recipient immune repertoire present at the time of BM transfer. These data show transfer of antigen-encoding BM under immune-preserving conditions modulates the established peripheral immune repertoire in addition to that which emerges as a consequence of BM transfer, to limit antigen-specific B-cell priming and cognate antibody production.

**Figure 2.**
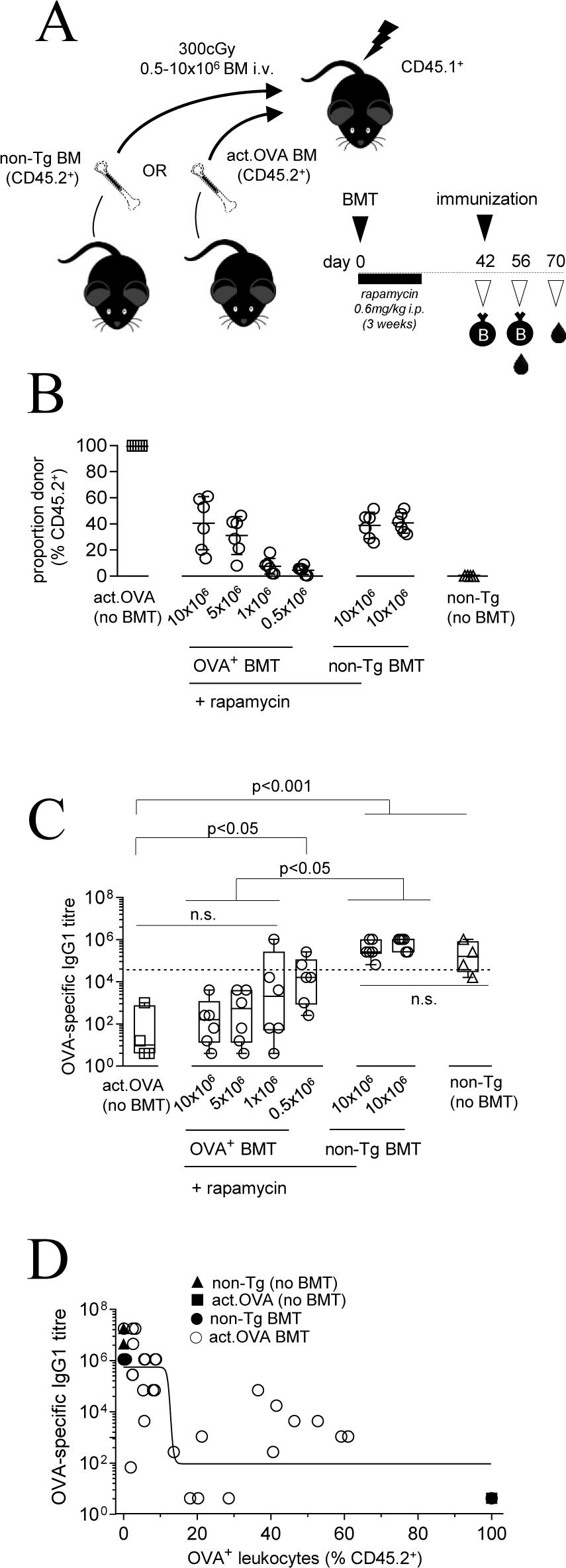
OVA-encoding bone marrow transfer under immune-preserving conditions limits OVA-specific antibody production in recipients. **A-D)** Graded doses of act.OVA BM (CD45.2^+^) or non-Tg BM (C57BL/6J, CD45.2^+^) were transferred to lightly-irradiated (300cGy) B6.SJL (CD45.1^+^) recipients. All mice receiving BM were treated with rapamycin (0.6mg/kg) with the exception of a control group receiving non-Tg BM. Untransplanted (act.OVA) and non-Tg (B6.SJL) mice were also used. 6 weeks after BM transfer, mice were immunized with OVA/CFA and 4 weeks later analyzed. **A)** Experimental layout. **B)** Chimerism was determined as the proportion of CD45.2^+^ cells in spleen 10 weeks post-transfer. **C)** OVA-specific IgG1 titres 4 weeks after OVA/CFA immunization. **D)** Relationship of OVA-specific IgG1 production to act.OVA chimerism. Plots show individual mice with mean±SD (B) or median and interquartile range (C). The dotted line indicates the antibody titre below which mice were considered ‘tolerant’. Data are pooled from 2 independent experiments. ANOVA with Tukey’s post-hoc test of log_10_-transformedtitres (C) and log-inhibitor vs response curve (D).

### Transfer of antigen-encoding BM under mild, immune preserving conditions purges antigen-specific B cells from the recipient peripheral B-cell repertoire

Having shown that transfer of antigen-encoding BM modulated cognate antibody production, we sought to understand whether B cells in recipients may have been directly modulated. We propose transfer of antigen-encoding BM leads to *de novo* expression of the antigen of interest in a form directly tolerogenic to B cells in recipients and one possible consequence of this might be deletion of B cells specific for the expressed antigen. Therefore, we established chimeras using low-dose irradiation and a dose of act.OVA BM that gave approximately 20% chimerism (Suppl. Fig. 3A). Six weeks after BM transfer we analyzed spleen and lymph nodes using a sensitive triple-tetramer assay that identifies naïve OVA-specific B cells [15] to determine the frequency of OVA-specific B cells. The overall frequency and number of mature (CD19^+^IgM^var^IgD^+^) B cell populations in spleen was similar between BM recipients and controls (Suppl. Fig. 3B–D) indicating similar B cell homeostasis across groups. Within the mature B cell compartment, OVA-specific B-cell frequency and number in spleen did not differ between non-Tg and act.OVA BM recipients (Fig. 3A). In contrast, in LN, OVA-specific B cells were reduced in both frequency and number in recipients of act.OVA BM relative to recipients of non-Tg BM (Fig. 3B). This replicates the pattern of OVA-specific B cell deletion seen in act.OVA mice relative to non-Tg controls (Fig.3A,B) and throughout our other extensive analyses of act.OVA mice. As the apparent deletion might have occurred within either the pre-existing recipient B cell population or that which developed from engrafted donor BM, we next examined each population. Within the recipient-derived LN B cell population, the number and frequency of OVA-specific mature LN B cells was reduced in recipients of act.OVA BM compared to recipients of non-Tg BM (Fig. 3C, Suppl. Fig. 3E). Although a similar trend was observed in the donor-derived B-cell population this was not statistically significant (Fig. 3C, Suppl. Fig. 3E). The frequency of OVA-specific B cells in LN of act.OVA BM recipients negatively correlated with OVA-expressing leukocyte chimerism (Fig. 3D, black circles) indicating deletion was tuned by the extent of OVA expression. Consistent with analyses of polyclonal self-reactive B cells (Taylor, Martinez et al. 2012) there was no difference in IgM expression (Fig. 3E) or tetramer binding (Fig. 3F) on those OVA-specific B cells that were present (likely representing those that escaped deletion) in the LN of OVA-encoding BM recipients. When considering the overall peripheral B cell populations (i.e. combining spleen and LN) our data suggest that approximately 35%of the overall OVA-specific B-cell pool was deleted.

**Figure 3.**
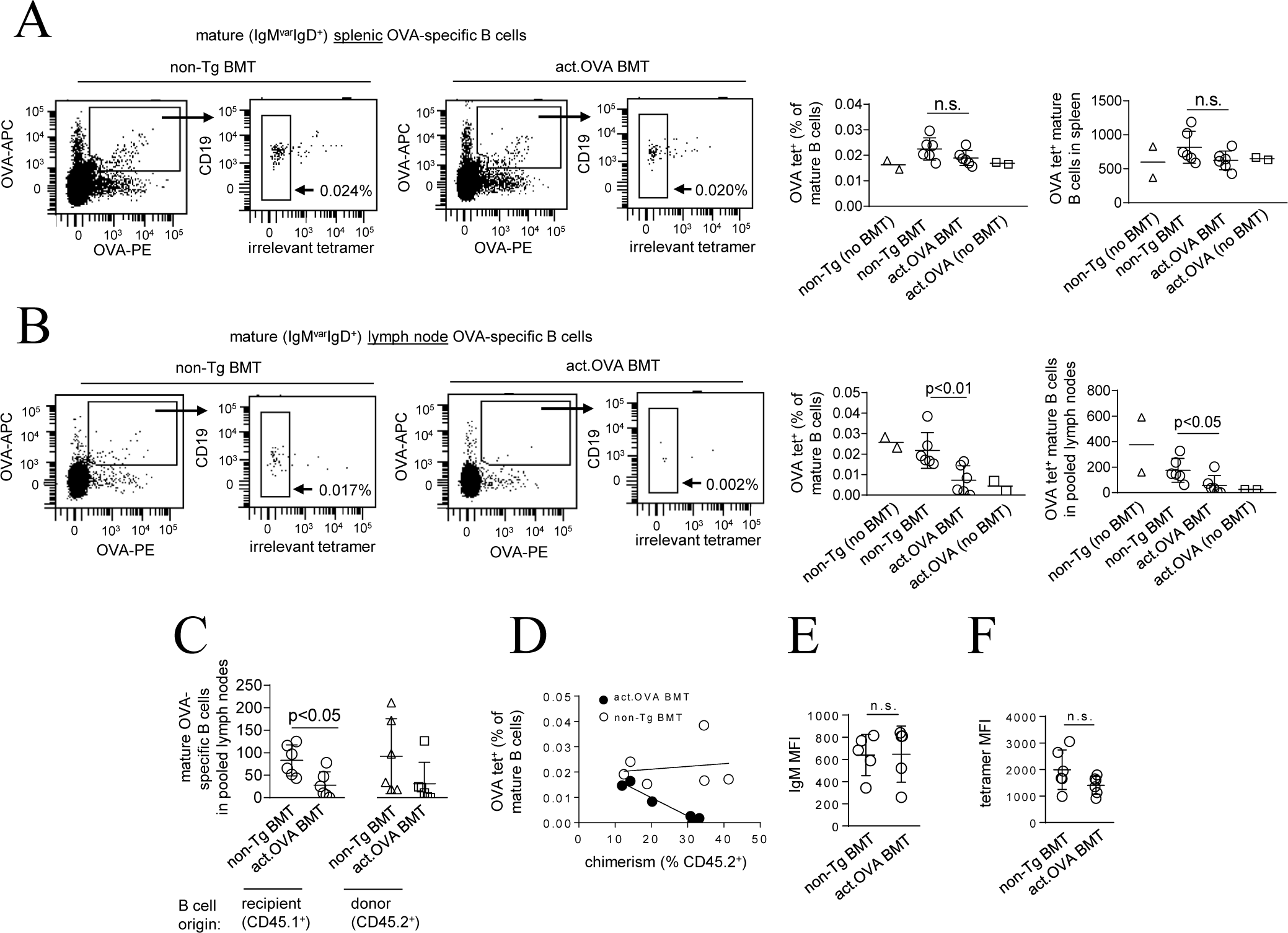
Recipient OVA-specific B cells are deleted following transfer of OVA-encoding BM under immune-preserving conditioning. **A-G**) 5×10^6^ act.OVA BM (CD45.2^+^) or non-Tg BM (C57BL/6J, CD45.2^+^) was transferred to lightly-irradiated (300cGy) B6.SJL (CD45.1^+^) recipients. All mice receiving BM were treated with rapamycin (0.6mg/kg). Untransplanted controls (act.OVA and non-Tg B6.SJL) were also used. Lymph nodes (pooled inguinal, axial, brachial, axillary) or spleens were analyzed 6 weeks after BM transfer. **A)** Representative FACS plots for discrimination of specific tetramer staining of mature (IgM^var^IgD^+^) B cells in spleen. Shown are cells pre-gated on mature B cells. Inset values represent OVA-specific B cells defined as OVA-tetramer^+ve^/irrelevant tetramer^−ve^ as a percentage of total mature B cells. Full gating strategy is available in Suppl. Fig. 3. Plots to the right show frequency and absolute number of mature splenic OVA-specific B cells. **B)** as in A, but for lymph node. **C)** Absolute number of mature OVA-specific B cells in LN of BM recipients defined by donor or recipient origin. **D)** Correlation between chimerism and mature OVA-specific B cell frequency in the lymph node. **E)** MFI of surface IgM on mature OVA-specific LN B cells. **F)** MFI of OVA-PE tetramer on mature OVA-specific LN B cells. Data are pooled from two independent experiments and plots show representative FACS plots or individual data points and mean±SD (A, B, C, E, F). Unpaired students *t*-test (A, B, C, E, F), linear regression (D).

### OVA-specific B cells are not recruited to germinal centres and fail to differentiate to plasmablasts in recipients of OVA-encoding BM

While deletion of OVA-specific B cells as a consequence of OVA-encoding BM transfer likely contributes to reduced OVA-specific antibody production, we sought to understand if this was the only contributing factor. Therefore, we immunized BM recipients and controls to define the B cell response in BM recipients and no BM transfer controls. We first examined differentiation of germinal centre (GC) B cells. The frequency and total overall number of all GC (CD19^+^GL7^+^CD38^−^) B cells in inguinal LN (iLN), draining the immunization site, was similar between recipients of act.OVA and non-Tg BM (Suppl. Fig. 4A–C) suggesting either no difference in the response to OVA or that the diverse antigenic makeup of CFA contributed to the overall response. Tetramer staining (Fig. 4A, Suppl. Fig. 4D) revealed that when the analysis was refined to OVA-specific B cells these were prominent among iLN GC B cells of non-Tg BM recipients after immunization, but not recipients of act.OVA BM (Fig. 4A, B, Suppl. Fig. 4D). The substantial population of OVA-specific GC B cell that developed in recipients of non-Tg BM were derived from B cells of both recipient and donor origin (Fig. 4C). In act.OVA BM, few OVA-specific GC B cells were identified of either recipient or donor origin (Fig. 4C), indicating that not only was the responsiveness of newly-developed donor-derived B cells ablated as might be expected, but that responsiveness within the pre-existing recipient B-cell pool was also ablated.

**Figure 4.**
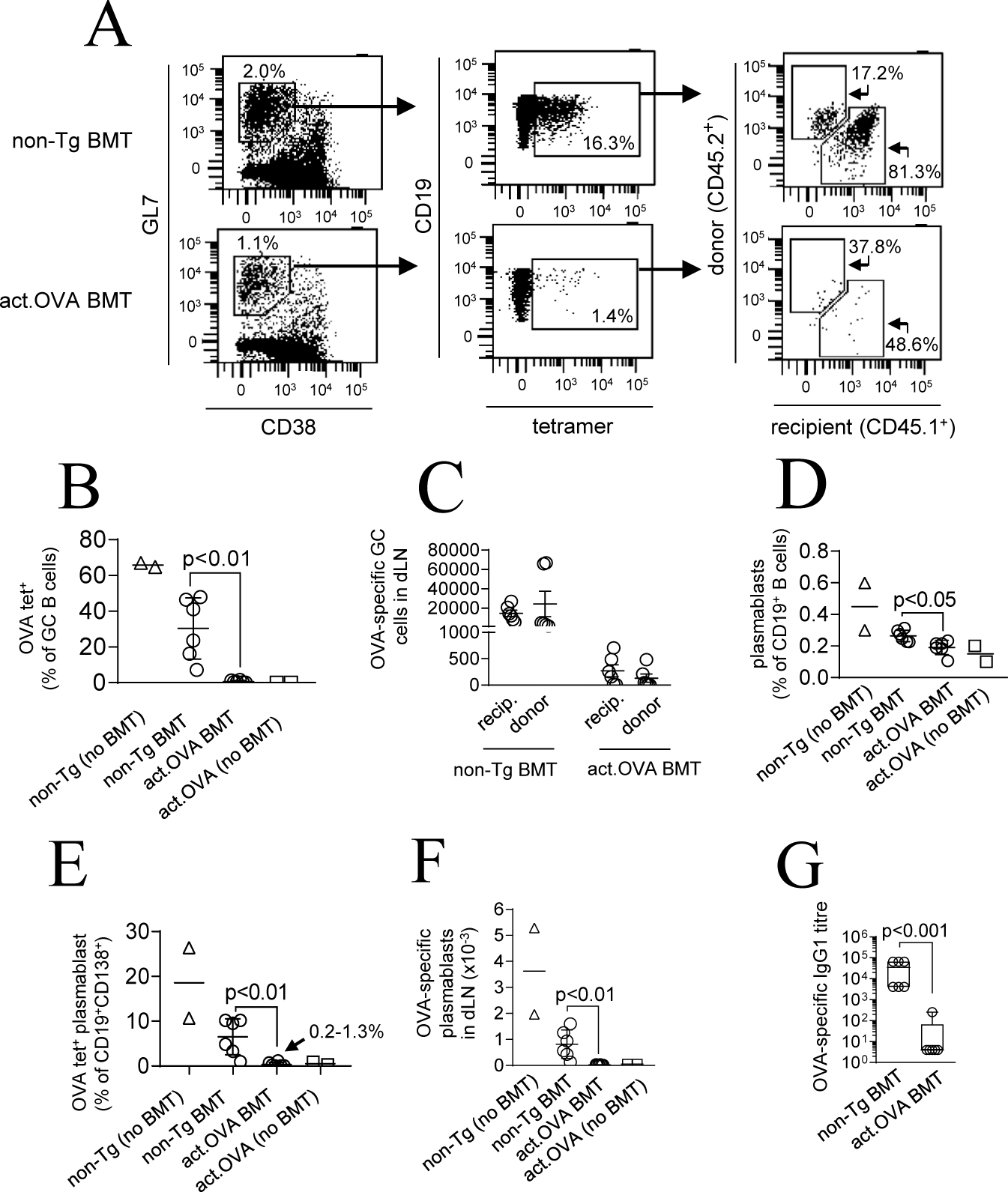
Germinal centre and plasmablast differentiation is restricted following OVA-encoding BM transfer. **A-H**) 5×10^6^ act.OVA BM (CD45.2^+^) or non-Tg BM (C57BL/6J, CD45.2^+^) was transferred to lightly-irradiated (300cGy) B6.SJL (CD45.1^+^) recipients. All mice receiving BM were treated with rapamycin (0.6mg/kg, 21 days). Untransplanted controls (act.OVA and non-Tg B6.SJL) were also used. Six weeks after BM transfer mice were euthanized and spleen, and pooled LN (brachial, axillary, inguinal, mesenteric) analyzed or immunized with OVA/CFA and two weeks later inguinal LN (left and right pooled) analyzed two weeks later. **A)** OVA-specific B cells were identified with OVA-PE tetramer. Representative gating for germinal centre (GL7^+^CD38^−^) cells (left panel), then OVA-tetramer binding (middle panel) and congenic CD45 expression (right panel). **B)** Frequency of OVA-specific GC B cells as a proportion of all GC B cells. **C)** Absolute number of OVA-specific GC B cells defined by donor or recipient origin in recipients of non-Tg or act.OVA BM. **D)** Frequency of plasmablasts (CD138^bright^) as a proportion of all CD19^+^ B cells. **E)** OVA-specific plasmablasts as a proportion of all plasmablasts. **G)** OVA-specific plasmablasts per pooled iLN. **F)** OVA-specific IgG1 titres two weeks post-immunization. Data are pooled from two independent experiments and show representative FACS plots or individual data points and mean±SD (B-F) or median with interquartile range. Unpaired student’s *t-*test (B, D-G) using log-transformed values for (G).

The reduced OVA-specific IgG1 titres in recipients of act.OVA BM 4 weeks after immunization (e.g. Fig 2C) suggested that OVA-specific plasmablast and/or plasma cell differentiation was also inhibited. When examined two weeks after immunization, the overall frequency (Fig. 4D) and total number of all plasmablasts (CD19^+^CD138^bright^) (Suppl. Fig. 4E) in iLN was reduced in recipients of actin.OVA compared to non-Tg BM recipients. Because plasmablasts retain some surface Ig we used OVA-tetramers to enumerate OVA-specific plasmablasts (Suppl. Fig. 4F). These comprised, on average, approximately 5% of the total plasmablast pool in recipients of non-Tg BM but were exceedingly rare (<1% of the total plasmablast pool) in recipients of act.OVA BM (Fig. 4E) and the total number of OVA-specific plasmablasts was minimal in recipients of act.OVA BM (Fig. 4F). In agreement with this, circulating OVA-specific IgG1 titres at this timepoint (2 weeks) after immunization were substantially reduced in OVA-encoding BM recipients relative to recipients of non-Tg BM (Fig. 4G). Together, these data show that antigen-encoding BM transfer not only leads to deletion of B cells specific for the expressed antigen, but also prevents/inhibits activation and passage through GC and plasma cell checkpoints, which here co-operate to limit antigen-specific antibody production from those antigen-specific B cells that escape deletion.

## Discussion

Current approaches to treating antibody-mediated pathologies manage the consequences of antibody-mediated effects using non-specific immune-suppression and/or attempt to indirectly control B cells, as the antibody source, through modulation of T cells. More recent approaches use biologics to broadly deplete B cells (e.g Rituximab) or block pathogenic antibodies (e.g. Omalizumab). Therapies that specifically prevent the development and/or ongoing production of deleterious antibodies by directly modulating antigen-specific B cells may be more effective. BM- or HSC-mediated gene therapy, used with the goal of expressing antigen(s) for tolerance induction, has been shown to ‘turn-off’ pathogenic T-cell responses (Coleman, Bridge et al. 2013). It is possible this could be extended to effectively, and antigen-specifically, control pathogenic antibody production. Studies using BCR transgenic mice and BM chimeras generated using immune-ablative approaches (Chung, Figgett et al. 2014), where all leukocytes are depleted, provide support for this concept. Although most studied (Yang, deGoma et al. 1998, Bracy and Iacomini 2000, Baranyi, Linhart et al. 2008, Chung, Figgett et al. 2014), immune-ablative approaches are not clinically desirable. Here we test a key advance for the field, use of mild conditions to facilitate engraftment of transferred HSC, which avoids immunoablation but generates an environment where B-cell specificities recognising the encoded antigen of interest, including those pre-existing at the time of BMT, would be deleted and/or modulated, thus demonstrating that mature B cells are subject to peripheral tolerance induced by BM transfer.

Development of BCR transgenic mice revolutionised study of B-cell dynamics. Early studies indicated a pre-eminent role for central B-cell tolerance mediated by deletion of antigen-specific B cells in the BM (Nemazee and Burki 1989, Hartley, Crosbie et al. 1991). Later, studies using more physiological models indicated that receptor editing was crucial for B-cell tolerance (Gay, Saunders et al. 1993, Tiegs, Russell et al. 1993) and that antigen-specific purging of B cells could occur in peripheral lymphoid organs (Russell, Dembić et al. 1991, Taylor, Martinez et al. 2012). The latter is important for therapeutic approaches aimed at stripping individual antigen specificities from within an established ‘mature’ B-cell repertoire and lends credence to approaches like the one tested here. A small number of studies have tested the principle that BM- or HSC-mediated gene therapy might modulate antigen-specific B cells but antigen-specific B cells have either not been tracked or, BCR transgenic B-cell models, which may not truly replicate the normal processes required for B cell tolerance, have been used (Chung, Figgett et al. 2014). For example, use of a myelin-oligodendrocyte glycoprotein-specific BCR transgenic system (Chung, Figgett et al. 2014) in an immune-ablative setting showed extensive deletion within the immature antigen-specific B-cell compartment which is consistent with early double transgenic models where receptor editing is limited (Hartley, Crosbie et al. 1991) but likely overestimates the role of central deletion. A key advance here is the use of multi-tetramer staining to define naïve antigen-specific B cells within a normal polyclonal repertoire with use of manipulated BCRs.

In the setting described here, we propose that BM transfer acts through peripheral mechanisms to control existing antigen-specific B cells present at the time of treatment, and then a combination of central and peripheral mechanisms control the repertoire of newly-developing B cells that arise from residual recipient hematopoiesis and that derived from the donor cells. To facilitate the testing of this hypothesis, we exploited a mild conditioning regime that enables engraftment of engineered BM but which largely preserves an existing immune repertoire (Coleman, Bridge et al. 2013, AL-Kouba, Wilkinson et al. 2017, Bhatt, Rudraraju et al. 2017). The challenge for study of B cells in a non-BCR-engineered setting has been the limited ability to detect naïve antigen-specific B cells within an unprimed B-cell repertoire. We recently validated a multi-tetramer approach for detection of rare naïve B cells in an unprimed repertoire (Brooks, Liu et al. 2018), and have used this to understand how antigen-encoding BM shapes the B-cell repertoire following engraftment of engineered BM. We also independently analyzed B cells emerging from transferred donor BM and those from the recipient leukocyte compartment from which we extrapolate changes that occur in the pre-existing B-cell repertoire that was present at the time of BM transfer. We found that in newly-emerging donor-derived B cells, antigen-specific B cells developed to maturity and were present in the spleen, but that a small fraction, and possibly those with highest affinity for OVA, were deleted in the periphery. This was prominent when LN were analyzed, suggesting deletion occurred at a very late stage of maturation. The extent of deletion found here is akin to that reported in studies of others in which the BCR repertoire is not directly genetically manipulated (Wardemann, Yurasov et al. 2003, Taylor, Martinez et al. 2012, Zikherman, Parameswaran et al. 2012). We also found a similar pattern of deletion in OVA-specific B cells of recipient origin, thereby providing evidence that the pre-existing B cell repertoire present at the time of BMT is modified by engineered BM transfer. This is an important finding as any therapeutic approach aimed at limiting deleterious antibody production should have the capacity to purge antigen responsiveness from an already established, pre-exiting B-cell repertoire. Although not tested here, it is possible that those emerging OVA-specific B cells that escaped deletion may have undergone receptor editing during development thereby limiting their BCR affinity for OVA.

In addition to mechanisms that alter diversity within the physical repertoire, such as deletion, we found that the responsiveness of OVA-specific B cells and OVA-specific antibody production was also limited. Anergy is enforced in B cells, by chronic BCR ligation (Cambier, Gauld et al. 2007). It might therefore be mooted that those OVA-specific B cells not deleted after OVA-encoding BM transfer, either within the pre-existing repertoire or that which develops subsequent to BM transfer, could be held in an anergic state by OVA expression consequent to BM transfer. Certainly, in both compartments, in recipients of OVA-encoding BM, very few OVA-specific B cells were recruited into GC reactions in response to OVA/CFA immunisation indicating a strong level of control at the GC checkpoint which regulates entry of B cells to a site where expansion and affinity maturation occurs. Interestingly, of the few OVA-specific B cells that had differentiated to a GC phenotype, virtually none progressed to become antibody-producing cells detected as OVA-tetramer^+ve^ plasmablasts, demonstrating that GC B cell and plasmablast differentiation of OVA-specific cells were both strongly inhibited and this together blocked antibody production. Of note, this occurred despite the significant differentiation of GC B cells and plasmablasts that occurred in parallel, that was presumably directed at immunogenic components of CFA such as mycobacterial antigens. The impaired OVA-specific antibody production is consistent with the suggestion that OVA-specific B cells were ‘anergised’ after OVA-encoding BM transfer, but could equally reflect a loss of OVA-specific T-cell responsiveness. As there is no reliable phenotypic means to determine whether B cells have been rendered anergic, and the few markers or phenotypic characteristics described appear useful only in BCR Tg settings, further studies are required to refine the mechanism of unresponsiveness.

It is generally considered that DC induce peripheral T-cell tolerance, but whether particular leukocytes are important for inducing peripheral B-cell tolerance is unclear, but follicular DC, which are not BM-derived, and therefore would not arise after BM transfer, may be involved. Indeed, targeting antigen expression to follicular DC induces developmental arrest at the transitional stage, and promotes apoptosis within antigen-specific B cells (Yau, Cato et al. 2013). It cannot be ruled out that antigen transfer to fDC contributes to the outcome here, but this does not limit efficacy of the approach. How targeting antigen expression to different cell types might influence the effectiveness of antigen-encoding BM transfer and, indeed, which tolerance mechanisms are active requires further study but it is likely that antigen-specific B-cell deletion and induction of unresponsiveness is dependent on chronic antigen recognition and so proximity of B cells to the antigen-expressing cells may be an important factor. In keeping with previous studies showing that widely-expressed and membrane-bound antigens appear to be most tolerogenic for B cells, our studies focussed on a setting where membrane-bound antigen would be expressed widely among the progeny of transferred BM. Notably, after BM transfer, antigen would be expressed solely by the progeny of transferred BM making it restricted to leukocytes, for example, within BM and peripheral lymphoid tissues in contrast to the donor act.OVA mice where all cells, including non-hematopoietic (e.g. stromal) cells, express antigen. Considering this, the data indicate that even relatively low proportions (e.g. ~10-20%) of antigen-expressing cells of haematopoietic origin are sufficient to induce both central and peripheral modulation of the B cell repertoire. This is consistent with a low abundance of antigen that is effectual in other settings (Adelstein, Pritchard-Briscoe et al. 1991, Gaudin, Hao et al. 2004, Tian, Bagley et al. 2006). Alternative promoters that target DC or even other leukocyte subsets may be equally effective or engineering neoantigen expression at distinct anatomical sites of autoimmune attack may also be effective at turning off B-cell responsiveness. The therapeutic value of defining a threshold for the antigen abundance or cell-specific requirement is that it can guide the tailoring of immune-preserving procedures so that they yield this level of antigen abundance to ensure therapeutic efficacy. A potentially key element, and one that might be restricted to gene-therapy approaches to immunotherapy is that antigen is successfully delivered in a form that is directly and highly tolergenic to B cells, viz., chronically-expressed with conformational fidelity and cell-surface anchored that cannot be replicated by ‘vaccine’-like’ approaches.

A challenge for therapy of deleterious antibody production, once established, is that it is maintained by long-lived plasma cells and plasma cells arising from activation of memory B cells. Plasma cells downregulate many cell surface proteins, including BCR, which render them impervious to agents that effectively deplete other B cell subsets (Hiepe and Radbruch 2016). Balancing this, newer agents may promote approaches to limiting the contribution of long-lived plasma cells to disease. Therefore, the ability to limit activation of newly–arisen naïve B cells and those existing within the repertoire, as shown here, or activation of memory B cells and their subsequent differentiation to plasma cells may be important to therapeutic outcomes. For example, tolerising T and B cells has been shown to halt or reverse disease progression in some preclinical models of antibody-mediated disease (Pozsgay, Szekanecz et al. 2017), lending credence to this suggestion. It does, however, remain to be directly determined whether responsiveness of memory B cells is limited by transfer of antigen-encoding BM. Alternatively, combination therapies might be leveraged to temporarily dampen plasma cell responses while inducing tolerance in pathogenic B cells.

We demonstrate that antigen-encoding BM transfer performed under conditions that preserve immunity in recipients induces robust deletion of antigen-specific B cells in both the repertoire present at the time of BM transfer and that which develops after BM transfer. Therefore, antigen-specific B cells within an established B-cell repertoire are susceptible to *de novo* tolerance induction and this can be achieved by transfer of gene-engineered bone marrow which leads to delivery of antigen in form that is directly tolerogenic to B cells. Furthermore, delivery of an antigen of interest using BM transfer ablates antigen-specific antibody production. The approach described here provides important proof-of-principle, interesting potential clinical opportunities and addresses some of the limitations of current approaches that act indirectly to modulate antigen-specific B-cell responsiveness. This adds further dimensions to the utility of antigen-encoding bone marrow transfer as an immunotherapeutic tool.

## Experimental procedures

### Mice

6-8 week-old C57BL/6J(Arc) (CD45.2^+^) and B6.SJL-Ptprc^a^Pep3^b^/BoyJ(Arc) (CD45.1^+^) mice were purchased from the Animal Resources Facility (Perth, Australia). Actin.OVA (act.OVA) mice (CD45.2^+^) express membrane-bound full-length ovalbumin (OVA) under the control of a β-actin promoter (Ehst, Ingulli et al. 2003). Mice were maintained under specific pathogen-free conditions at the TRI Biological Resources Facility (Brisbane, Australia) and typically used at 8-16 weeks of age. Studies were approved by the University of Queensland Ethics Committee.

### Bone marrow transplantation

Bone marrow (BM) transplants were performed as described (Bhatt, Rudraraju et al. 2017). Briefly, femurs and tibiae from donor mice were harvested and BM flushed with cold PBS/FCS (2.5% v/v). BM was passaged through a 21g needle to prepare a single cell suspension and erythrocytes lysed using ammonium chloride buffer. BM was resuspended in mouse-tonicity phosphate buffered saline (MT-PBS) for transfer. For myeloablative (1100cGy) procedures, mice received fractionated irradiation (2 x 550cGy) three hours apart, 4 hours prior to transplant and neomycin (1g/L) in drinking water for 3 weeks post-transplant. For non-myeloablative (300cGy) procedures, mice received a single dose of irradiation. Some mice were administered rapamycin (0.6mg/kg) daily for 3 weeks post-transplant (Bhatt, Rudraraju et al. 2017).

### Immunizations

For immunization of act.OVA chimeras, OVA (Sigma, Grade V, A5503) 2mg/mL in PBS was emulsified in Complete Freund’s adjuvant (CFA, Sigma F5881 or Difco 23810 – MO, USA) using a mini-BeadBeater (Biospec Products – OK, USA) and 100µl injected s.c. at the tail base.

### Flow cytometry

OVA-labelled tetramers have been described (Brooks, Liu et al. 2018). OVA and HEL were biotinylated in-house and tetramerised with streptavidin-PE (Biolegend – CA, USA), streptavidin-APC (Biolegend – CA, USA) or streptavidin-BV510 (BD Bioscience – CA, USA) for two hours on ice. Tetramers were diluted 1:200 (19.2nM) for staining. IgM-PECy7 (RMM-1), IgD-BUV395 (11-26c.2a), CD4-FITC (GK1.5) or CD4-PE (RM4-4), CD8-PerCPCy5.5 (53-6.7) or CD8-FITC (53-6.7) or CD8-PECy7 (53-6.7), CD11c-FITC (N418) or CD11c-PE (N418), B220-APC (RA3-B62), CD19-APC (6D5) or CD19-BV785 (6D5), CD45.1-APC (A20) or CD45.1-FITC (A20), CD45.2-FITC (104) or CD45.2-PE-Cy7 (104), and GL7-Pacific Blue (GL7) were purchased from Biolegend (CA, USA). CD138-AF700 (281-2), CD45.1-BV711 (A20), CD45.2-BUV737 (104), IgG1-biotin (A85-1), and IgG2a,2b-biotin (R2-40) were purchased from BD Bioscience (CA, USA). NK1.1 (PK136), Gr1 (RB6-8C5) and CD11b (M1-70) were purchased from BioXCell (NH, USA) or grown in-house and labelled with FITC in house.

Pooled lymph nodes (inguinal, brachial, axillary, mesenteric) and spleens for naïve mice, or inguinal lymph nodes for immunized mice, were collected and pressed through a 70µM strainer (Corning – NY, USA) to create single cell suspensions. Erythrocytes were lysed (spleens) using ammonium chloride buffer. Antibody and tetramer staining was as described (Brooks, Liu et al. 2018) and all staining steps were performed on ice. Absolute cell number was enumerated using a bead-based counting assay with Flow-Count^TM^ Fluorospheres (Beckman Coulter – CA, USA) as described (Brooks, Liu et al. 2018). Data were collected on an LSR Fortessa X-20 (BD Bioscience – CA, USA) and analyzed using DIVA version 8 (BD Bioscience – CA, USA).

### ELISA

For OVA-specific IgG1 ELISA, plasma was serially diluted (4-fold dilutions) and added to ELISA plates (Maxisorp, Nunc – MA, USA) pre-coated with 10µg/mL OVA (Sigma Grade V, A5503 – MO, USA). OVA-specific IgG1 was detected using biotinylated anti-mouse IgG1 monoclonal antibody (0.02µg/mL RMG1-1, Biolegend – CA, USA) followed by SA-HRP (1:2000 dilution #P030701-2, DAKO – CA, USA) and visualised with TMB (#421101 Biolegend – CA, USA). Absorbance was measured at 450nm and corrected for absorbance at 540nm using a MultiSkan Go spectrophotometer (Thermofisher – MA, USA).

### Statistical analysis

OVA-specific IgG1 titre plots show untransformed data, statistical testing of antibody titres was performed on log_10_-transformed titres. Dose/inhibition curves were calculated using GraphPad Prism V7 (GraphPad Software, Inc. – CA, USA). Comparison of means used student t-test or ANOVA with Tukey post-hoc analysis for multiple comparisons (GraphPad Prism V8).

## Author contributions

Conceptualization: JFB and RJS; Methodology: JFB and RJS; Investigation JFB and RJS; Writing –Original Draft: JFB and RJS; Writing –Reviewing and Editing: JFB, RJS, JMD and JWW. Supervision JMD and JWW. All authors have approved the manuscript.

## Acknowledgements

JFB was supported by a Research Training Program Scholarship and Children’s Hospital Foundation Top-up award (#50209, RPCPHD0072017), JMD by Queensland University of Technology, JWW by a Fellowship from Perpetual Trustees and RJS by a UQ Vice Chancellor’s Senior Research Fellowship. The authors thank Associate Professor Julie Zikherman for critical reading of the manuscript and the TRI Flow Cytometry Core Facility for excellent flow cytometry support.

---

**Figure S1.**
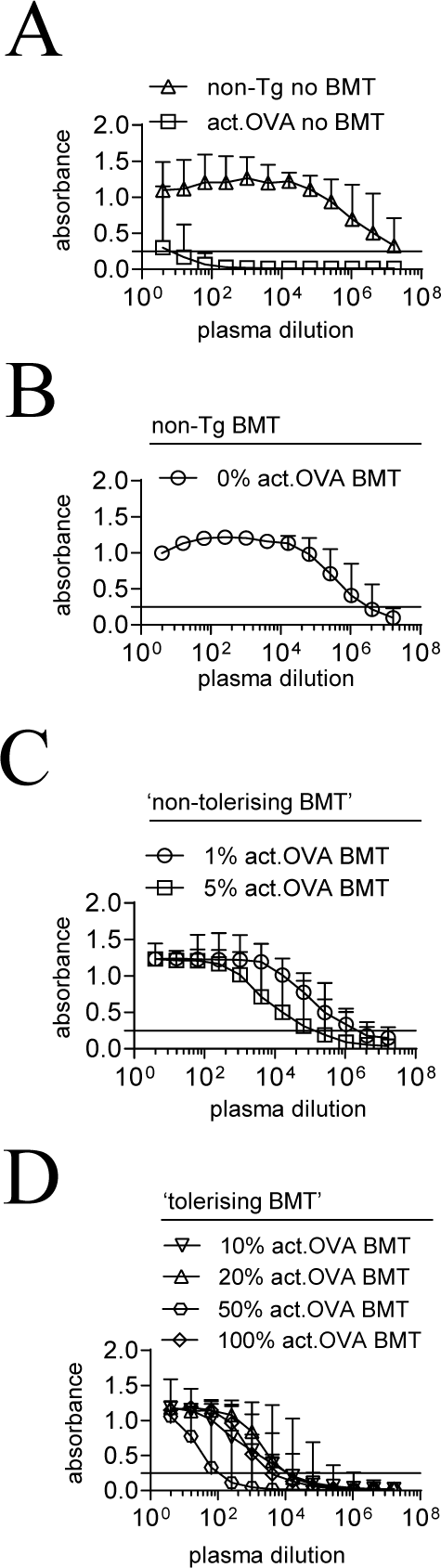
Production of OVA-specific IgG1 in BM chimeras and controls. **A-D)** Mixtures of OVA-encoding (CD45.2^+^) and non-Tg (CD45.1^+^) BM were transferred to irradiated (1100cGy) recipients (CD45.2^+^). BM recipients and controls were immunized with OVA/CFA 6 weeks after BM transfer and analysed 28 days later. **A)** Titration curves for OVA-specific IgG1 from non-Tg and act.OVA (no BMT) controls. **B)** Titration curves for non-Tg BMT recipients. **C)** Titration curves for 1% and 5% act.OVA BM chimeras. **D)** Titration curves for 10-100% act.OVA BM chimeras. Data are shown as median± interquartile range. Data are pooled from two independent experiments. ‘non-tolerising’ BMT refers to recipients of act.OVA BM where titres did not differ significantly from non-Tg (no BMT) controls. ‘tolerising’ BMT refers to recipients of act.OVA BM where titres were significantly reduced from non-Tg (no BMT) controls (taken from Figure 1C).

**Figure S2.**
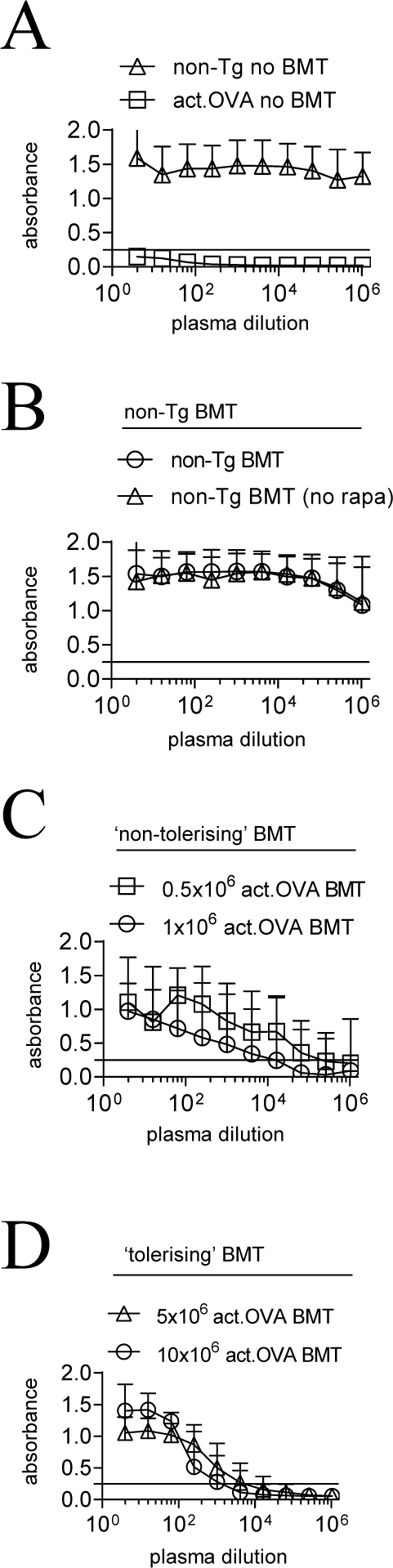
Production of OVA-specific IgG1 in controls and after after BM transfer using low-dose irradiation. **A-D)** Increasing doses of act.OVA (CD45.2^+^) BM (0.5-10×10^6^ cells per mouse) or non-Tg (C57BL/6J) BM (CD45.2^+^) BM (10×10^6^ cells per mouse only) were transferred to lightly-irradiated (300cGy) CD45.1^+^ B6.SJL recipients. Low-dose rapamycin (0.6mg/kg) was administered for 3 weeks posttransplant as indicated. BM recipients and controls were immunized with OVA/CFA 6 weeks after BM transfer and analysed 28 days later. **A)** Titration curves for OVA-specific IgG1 from non-Tg and act.OVA (no BMT) controls. **B)** Titration curves for non-Tg BMT recipients. **C)** Titration curves for 0.5×10^6^ and 1×10^6^ act.OVA BM chimeras. **D)** Titration curves for 5×10^6^ and 10×10^6^ act.OVA BM chimeras. Data are shown as median± interquartile range. Data are pooled from two independent experiments. ‘non-tolerising’ BMT refers to recipients of act.OVA BM where titres did not differ significantly from non-Tg (no BMT) controls. ‘tolerising’ BMT refers to recipients of act.OVA BM where titres were significantly reduced from non-Tg (no BMT) controls (taken from Figure 2C).

**Figure S3.**
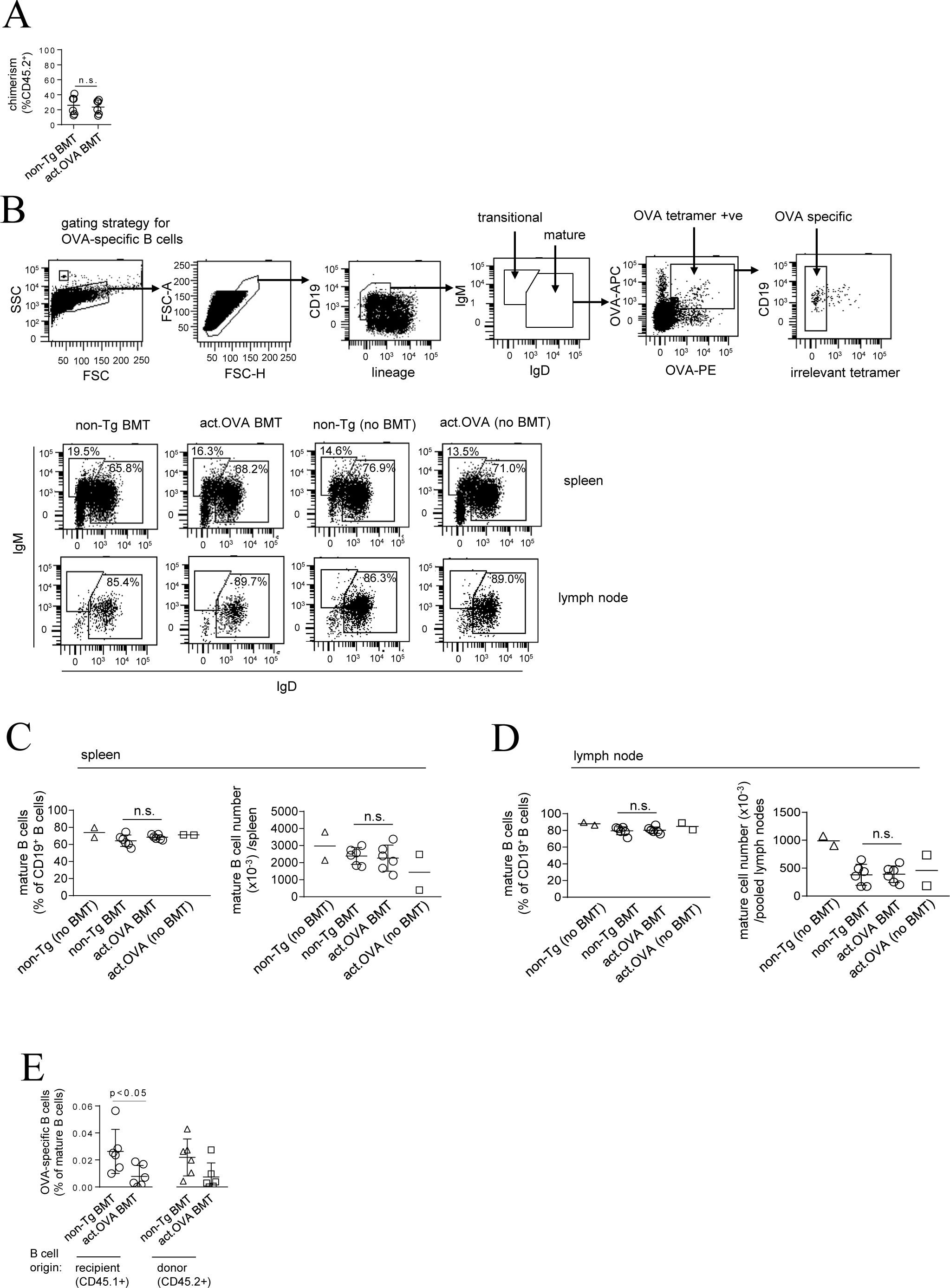
Analysis of naïve B cell repertoire following BM transfer using low-dose irradiation. **A-E)** 5×10^6^ act.OVA (CD45.2^+^) or non-Tg (C57BL/6) BM (CD45.2^+^) was transferred to lightly-irradiated (300cGy) recipients (CD45.1^+^). Mice received low dose rapamycin (0.6mg/kg) for 3 weeks post-transplant. Mice were analysed 6 weeks post-transfer. **A)** Engraftment levels of donor CD45.2^+^ BM in spleen at week 6 after BMT. **B)** Spleen and lymph nodes were harvested and stained and analysed as indicated and with OVA-tetramers to identify OVA-specific B cells. Analysis of transitional and mature B cells in spleen and mature B cells in lymph nodes, gated as shown. Inset values represent proportion gated as a percentage of CD19^+^ B cells. Mature OVA-specific B cells were analysed using the strategy shown. **C)** Percentage and number of splenic mature (IgM^var^IgD^+^) B cells. **D)** Percentage and number of lymph node mature (IgM^var^IgD^+^) B cells. **E)** OVA-specific B cells dissected based on donor (CD45.2^+^) or recipient (CD45.1^+^) origin, represented as the percentage of mature (IgM^var^IgD^+^) B cells in the lymph node. Data are pooled from two independent experiments, each dot represents an individual mouse with mean±SD shown. Unpaired t-test (**A, C-E)**.

**Figure S4.**
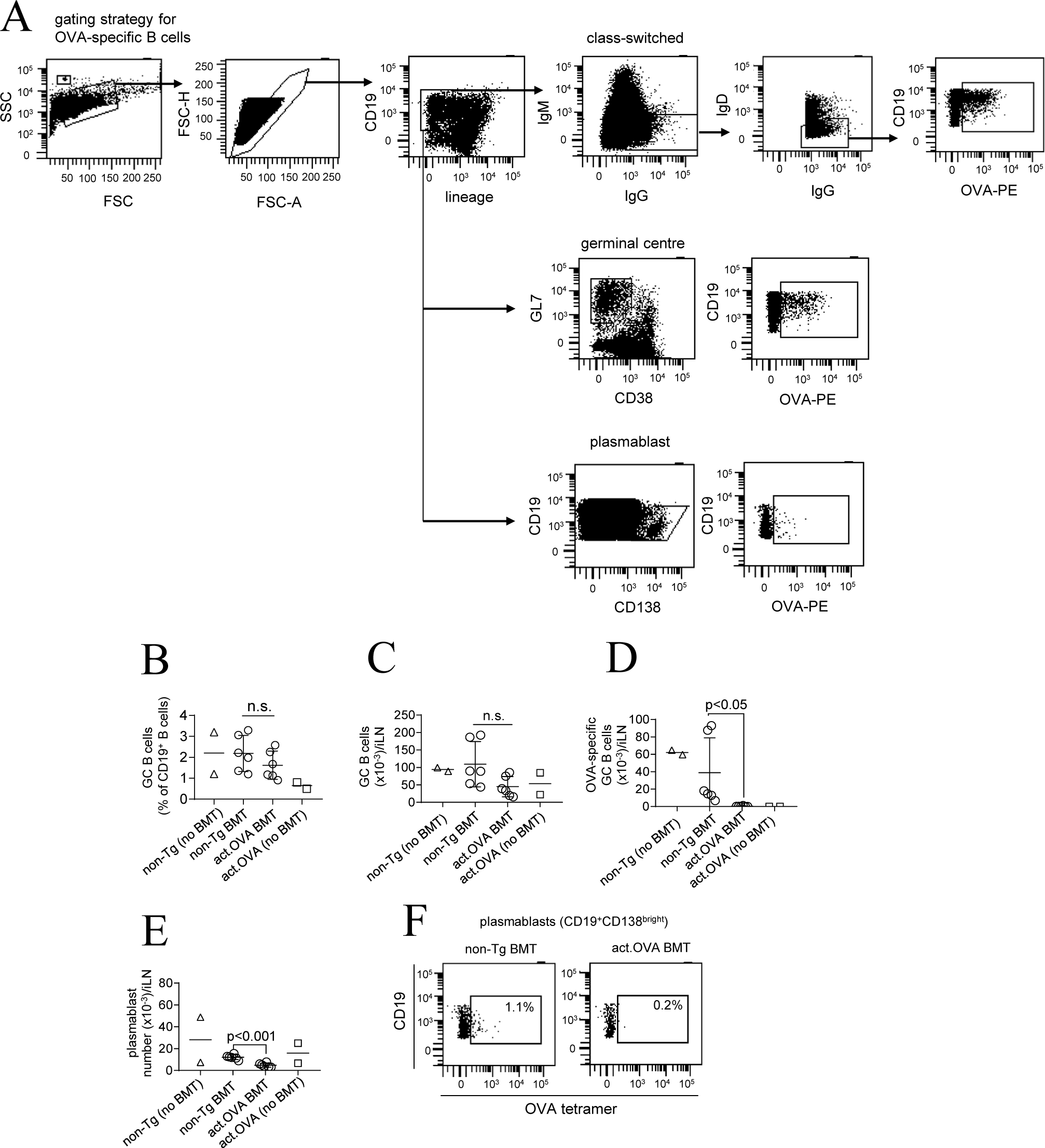
Analysis of germinal centre B cells and plasmablasts after immunization of non-myeloablative chimeras. **A-F)** 5×10^6^ act.OVA (CD45.2^+^) or non-Tg (C57BL/6) BM (CD45.2^+^) was transferred to lightly-irradiated (300cGy) recipients (CD45.1^+^). Mice received low dose rapamycin (0.6mg/kg) for 3 weeks post-transplant. Mice were immunized (OVA/CFA) 6 weeks posttransfer and analysed 4 weeks later.14 days after immunization, the draining (inguinal) lymph nodes were analysed. **A)** Representative gating strategy. **B, C)** Total number and percentage of GC B cells in the lymph nodes. GC B cells are represented as a proportion of CD19^+^ B cells. **D)** Number of OVA-specific GC B cells. **E)** Total number of plasmablasts. **F)** Representative tetramer staining of CD19^+^CD138^bright^ plasmablasts. Data are representative FACS plots or individual data points pooled from two independent experiments. Bars denote mean±SD. Unpaired t-test **(B-E)**.

## References

Adelstein, S., H. Pritchard-Briscoe, T. A. Anderson, J. Crosbie, G. Gammon, R. H. Loblay, A. Basten and C. C. Goodnow (1991). “Induction of self-tolerance in T cells but not B cells of transgenic mice expressing little self antigen.” Science 251(4998): 1223–1225.

AL-Kouba, J., A. Wilkinson, M. R. Starkey, R. Rudraraju, R. Werder, S.-C. Law, T. Liu, J. C. Horvat, J. F. Brooks, G. R. Hill, J. M. Davies, S. Phipps, P. M. Hansbro and R. J. Steptoe (2017). “Allergen-encoding bone-marrow transfer inactivates allergic T-cell responses, alleviating airways inflammation.” JCI Insight 2(11): e85742.

Baranyi, U., B. Linhart, N. Pilat, M. Gattringer, J. Bagley, F. Muehlbacher, J. Iacomini, R. Valenta and T. Wekerle (2008). “Tolerization of a type I allergic immune response through transplantation of genetically modified hematopoietic stem cells.” J Immunol 180(12): 8168–8175.

Bhatt, K. H., R. Rudraraju, J. F. Brooks, J.-W. Jun, R. Galea, J. W. Wells and R. J. Steptoe (2017). “Short-course rapamycin treatment enables engraftment of immunogenic gene-engineered bone marrow under low-dose irradiation to permit long-term immunological tolerance.” Stem Cell Res Ther 8(1): 57.

Bracy, J. L. and J. Iacomini (2000). “Induction of B-cell tolerance by retroviral gene therapy.” Blood 96(9): 3008–3015.

Brooks, J. F., X. Liu, J. M. Davies, J. W. Wells and R. J. Steptoe (2018). “Tetramer-based identification of naïve antigen-specific B cells within a polyclonal repertoire.” European Journal of Immunology 48(7): 1251–1254.

Cambier, J. C., S. B. Gauld, K. T. Merrell and B. J. Vilen (2007). “B-cell anergy: from transgenic models to naturally occurring anergic B cells?” Nat Rev Immunol 7(8): 633–643.

Chung, J. Y., W. Figgett, K. Fairfax, C. Bernard, J. Chan, B. H. Toh, F. Mackay and F. Alderuccio (2014). “Gene therapy delivery of myelin oligodendrocyte glycoprotein (MOG) via hematopoietic stem cell transfer induces MOG-specific B cell deletion.” J Immunol 192(6): 2593–2601.

Cogollo, E., M. A. Silva and D. Isenberg (2015). “Profile of atacicept and its potential in the treatment of systemic lupus erythematosus.” Drug Des Devel Ther 9: 1331–1339.

Coleman, M. A., J. A. Bridge, S. W. Lane, C. M. Dixon, G. R. Hill, J. W. Wells, R. Thomas and R. J. Steptoe (2013). “Tolerance induction with gene-modified stem cells and immune-preserving conditioning in primed mice: restricting antigen to differentiated antigen-presenting cells permits efficacy.” Blood 121(6): 1049–1058.

Coleman, M. A., C. F. Jessup, J. A. Bridge, N. H. Overgaard, D. Penko, S. Walters, D. J. Borg, R. Galea, J. M. Forbes, R. Thomas, P. T. H. Coates, S. T. Grey, J. W. Wells and R. J. Steptoe (2016). “Antigen-encoding bone marrow terminates islet-directed memory CD8+ T-cell responses to alleviate islet transplant rejection.” Diabetes 65(5): 1328–1340.

Ehst, B. D., E. Ingulli and M. K. Jenkins (2003). “Development of a Novel Transgenic Mouse for the Study of Interactions Between CD4 and CD8 T Cells During Graft Rejection.” Am J Transplant 3(11): 1355–1362.

Gaudin, E., Y. Hao, M. M. Rosado, R. Chaby, R. Girard and A. A. Freitas (2004). “Positive selection of B cells expressing low densities of self-reactive BCRs.” J Exp Med 199(6): 843–853.

Gay, D., T. Saunders, S. Camper and M. Weigert (1993). “Receptor editing: an approach by autoreactive B cells to escape tolerance.” J Exp Med 177(4): 999–1008.

Getahun, A. and J. C. Cambier (2019). “Non-Antibody-Secreting Functions of B Cells and Their Contribution to Autoimmune Disease.” Annu Rev Cell Dev Biol.

Goodnow, C. C., J. Crosbie, S. Adelstein, T. B. Lavoie, S. J. Smith-Gill, R. A. Brink, H. Pritchard-Briscoe, J. S. Wotherspoon, R. H. Loblay, K. Raphael, R. J. Trent and A. Basten (1988). “Altered immunoglobulin expression and functional silencing of self-reactive B lymphocytes in transgenic mice.” Nature 334(6184): 676–682.

Hale, M., D. J. Rawlings and S. W. Jackson (2018). “The long and the short of it: insights into the cellular source of autoantibodies as revealed by B cell depletion therapy.” Curr Opin Immunol 55: 81–88.

Hartley, S. B., J. Crosbie, R. Brink, A. B. Kantor, A. Basten and C. C. Goodnow (1991). “Elimination from peripheral lymphoid tissues of self-reactive B lymphocytes recognizing membrane-bound antigens.” Nature 353(6346): 765–769.

Hartley, S. B., J. Crosbie, R. Brink, A. B. Kantor, A. Basten and C. C. Goodnow (1991). “Elimination from peripheral lymphoid tissues of self-reactive B lymphocytes recognizing membrane-bound antigens.” Nature 353: 765.

Hiepe, F. and A. Radbruch (2016). “Plasma cells as an innovative target in autoimmune disease with renal manifestations.” Nat Rev Nephrol 12(4): 232–240.

Jelcic, I., F. Al Nimer, J. Wang, V. Lentsch, R. Planas, I. Jelcic, A. Madjovski, S. Ruhrmann, W. Faigle, K. Frauenknecht, C. Pinilla, R. Santos, C. Hammer, Y. Ortiz, L. Opitz, H. Gronlund, G. Rogler, O. Boyman, R. Reynolds, A. Lutterotti, M. Khademi, T. Olsson, F. Piehl, M. Sospedra and R. Martin (2018). “Memory B Cells Activate Brain-Homing, Autoreactive CD4(+) T Cells in Multiple Sclerosis.” Cell 175(1): 85–100.e123.

Kerkman, P. F., Y. Rombouts, E. I. van der Voort, L. A. Trouw, T. W. Huizinga, R. E. Toes and H. U. Scherer (2013). “Circulating plasmablasts/plasma cells as a source of anticitrullinated protein antibodies in patients with rheumatoid arthritis.” Ann Rheum Dis 72(7): 1259–1263.

Marino, E., S. N. Walters, J. E. Villanueva, J. L. Richards, C. R. Mackay and S. T. Grey (2014). “BAFF regulates activation of self-reactive T cells through B-cell dependent mechanisms and mediates protection in NOD mice.” Eur J Immunol 44(4): 983–993.

Nemazee, D. A. and K. Burki (1989). “Clonal deletion of B lymphocytes in a transgenic mouse bearing anti-MHC class I antibody genes.” Nature 337(6207): 562–566.

Neubert, K., S. Meister, K. Moser, F. Weisel, D. Maseda, K. Amann, C. Wiethe, T. H. Winkler, J. R. Kalden, R. A. Manz and R. E. Voll (2008). “The proteasome inhibitor bortezomib depletes plasma cells and protects mice with lupus-like disease from nephritis.” Nat Med 14(7): 748–755.

Pozsgay, J., Z. Szekanecz and G. Sarmay (2017). “Antigen-specific immunotherapies in rheumatic diseases.” Nat Rev Rheumatol 13(9): 525–537.

Ramanujam, M., X. Wang, W. Huang, Z. Liu, L. Schiffer, H. Tao, D. Frank, J. Rice, B. Diamond, K. O. Yu, S. Porcelli and A. Davidson (2006). “Similarities and differences between selective and nonselective BAFF blockade in murine SLE.” J Clin Invest 116(3): 724–734.

Russell, D. M., Z. Dembić, G. Morahan, J. F. Miller, K. Bürki and D. Nemazee (1991). “Peripheral deletion of self-reactive B cells.” Nature 354(6351): 308–311.

Sabatos-Peyton, C. A., J. Verhagen and D. C. Wraith (2010). “Antigen-specific immunotherapy of autoimmune and allergic diseases.” Current Opinion in Immunology 22(5): 609–615.

Stathopoulos, P., A. Kumar, R. J. Nowak and K. C. O’Connor (2017). “Autoantibody-producing plasmablasts after B cell depletion identified in muscle-specific kinase myasthenia gravis.” JCI Insight 2(17).

Steptoe, R. J., J. M. Ritchie and L. C. Harrison (2003). “Transfer of hematopoietic stem cells encoding autoantigen prevents autoimmune diabetes.” J Clin Invest 111(9): 1357–1363.

Suurmond, J. and B. Diamond (2015). “Autoantibodies in systemic autoimmune diseases: specificity and pathogenicity.” J Clin Invest 125(6): 2194–2202.

Taylor, J. J., R. J. Martinez, P. J. Titcombe, L. O. Barsness, S. R. Thomas, N. Zhang, S. D. Katzman, M. K. Jenkins and D. L. Mueller (2012). “Deletion and anergy of polyclonal B cells specific for ubiquitous membrane-bound self-antigen.” J Exp Med 209(11): 2065–2077.

Teng, Y. K., G. Wheater, V. E. Hogan, P. Stocks, E. W. Levarht, T. W. Huizinga, R. E. Toes and J. M. van Laar (2012). “Induction of long-term B-cell depletion in refractory rheumatoid arthritis patients preferentially affects autoreactive more than protective humoral immunity.” Arthritis Res Ther 14(2): R57.

Tian, C., J. Bagley and J. Iacomini (2006). “Persistence of antigen is required to maintain transplantation tolerance induced by genetic modification of bone marrow stem cells.” Am J Transplant 6(9): 2202–2207.

Tiegs, S. L., D. M. Russell and D. Nemazee (1993). “Receptor editing in self-reactive bone marrow B cells.” J Exp Med 177(4): 1009.

Wardemann, H., S. Yurasov, A. Schaefer, J. W. Young, E. Meffre and M. C. Nussenzweig (2003). “Predominant autoantibody production by early human B cell precursors.” Science 301(5638): 1374–1377.

Wu, L. C. and H. Scheerens (2014). “Targeting IgE production in mice and humans.” Curr Opin Immunol 31: 8–15.

Yang, Y. G., E. deGoma, H. Ohdan, J. L. Bracy, Y. Xu, J. Iacomini, A. D. Thall and M. Sykes (1998). “Tolerization of anti-Galalpha1-3Gal natural antibody-forming B cells by induction of mixed chimerism.” J Exp Med 187(8): 1335–1342.

Yau, I. W., M. H. Cato, J. Jellusova, T. Hurtado de Mendoza, R. Brink and R. C. Rickert (2013). “Censoring of self-reactive B cells by follicular dendritic cell-displayed self-antigen.” J Immunol 191(3): 1082–1090.

Zikherman, J., R. Parameswaran and A. Weiss (2012). “Endogenous antigen tunes the responsiveness of naive B cells but not T cells.” Nature 489: 160.

